# Ancient DNA from a 2,700-year-old goitered gazelle (*Gazella subgutturosa*) confirms gazelle hunting in Iron Age Central Asia

**DOI:** 10.1101/2021.12.08.471591

**Authors:** André Elias Rodrigues Soares, Nikolaus Boroffka, Oskar Schröder, Leonid Sverchkov, Norbert Benecke, Torsten Günther

## Abstract

Central Asia has been an important region connecting the different parts of Eurasia throughout history and prehistory, with large states developing in this region during the Iron Age. Archaeogenomics is a powerful addition to the zooarchaeological toolkit for understanding the relation of these societies to animals. Here, we present the genetic identification of a goitered gazelle specimen (*Gazella subgutturosa*) at the site Gazimulla-Tepa, in modern-day Uzbekistan, confirming hunting of the species in the region during the Iron Age. The sample was directly radiocarbon dated to 2724-2439 calBP. A phylogenetic analysis of the mitochondrial genome places the individual into the modern variation of *G. subgutturosa*. Our data does represent both the first ancient DNA and the first nuclear DNA sequences of this species. The lack of genomic resources available for this gazelle and related species prevented us from performing a more in-depth analysis of the nuclear sequences generated. Therefore, we are making our sequence data available to the research community to facilitate other research of this nowadays threatened species which has been subject to human hunting for several millennia across its entire range on the Asian continent.

## Introduction

The Surkhandary Region in South-eastern Uzbekistan as part of the ancient region of Bactria has been a focus for archaeological research for many years. Belonging to the Achaemenid Empire, the “First Persian Empire”, it was later conquered by Alexander the Great. Excavations at the fortress of Kurganzol produced well-stratified samples of many faunal remains, architectural features and finds, mainly from the late 4th century BC, providing insights into Alexander’s campaigns in Central Asia [1]. Prior to Alexander’s arrival, several important settlements in the region were located close to the modern town of Bandykhan/Bandixon, near the southern edge of the Baysun Basin [1,2,3]. Among the sites in this region is Gazimulla-Tepa (Bandykhan III, Figure 1a), a city-type settlement around a citadel dating to the pre-Achaemenian period (Yaz IIB period), from the 9th to the 6th centuries BC [4,5]. The radiocarbon date falls into the “Hallstatt Plateau”, and therefore does not allow a precise calibrated date. However, the archaeological context is the pre-Achaemenian period, locally named Yaz IIB [3,6], so that a date before the middle of the 6^th^ century BC can be established. It was only under Cyrus II (the Great), that Central Asia was conquered and integrated into the Achaemenian Empire. This occurred around 546-540 BC [7], so that the site of Gazimulla-Tepe must be placed before the mid-6^th^ century BC.

**Figure 1:**
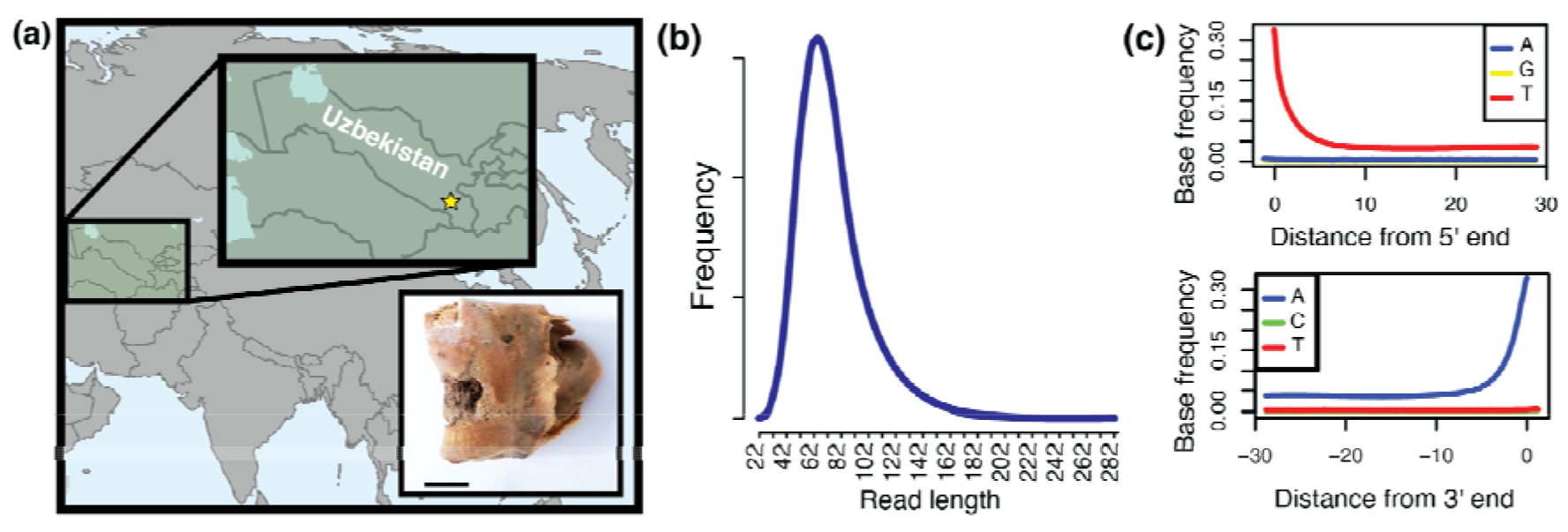
(a) Locality in which the bone fragment was found is indicated on the map with a yellow star. The insert contains a photo of the bone fragment (sample AGAZ005, Gazimullah Y:2006 F:3). Black bar indicates 1 cm length. (b) DNA fragment length distribution short sequences characteristic of ancient DNA. (c) Misincorporation plots indicating DNA damage patterns towards the end of the sequencing reads, also congruent with authentic ancient DNA fragments.

People living in this region largely relied on domesticated animal species as most faunal remains of that time in the region belong to species such as sheep, cattle and horses. Hunting still played a non-negligible role in their lives as a number of bones from wild mammals such as deer, gazelles and wild boars are found at the sites [8].

Ancient DNA approaches are a powerful addition to the zooarchaeological toolkit allowing to assign species, ancestry, biological sex and/or certain traits to faunal remains (e.g. [9,10]). However, most studies have focused on domesticated animals instead of wild species that were hunted. In this study we sequenced DNA from an approximately 2700 years old bovid bone fragment excavated at Gazimulla-Tepa during the year 2006 (Figure 1a). The fragment was found among mostly domestic animal remains but we identify it as belonging to a goitered gazelle (*Gazella subgutturosa*). The goitered gazelle was among the most commonly hunted animals in prehistoric central and western Asia [8,11–13], and yet this is the first ancient DNA generated from the species allowing us to genetically categorize its relationships with modern gazelle populations.

## Material and Methods

### DNA Extraction and Sequencing

We sampled ∼150 mg of bone fragment from the humerus bone specimen (Collection Accession ID Gazimullah Y:2006 F:3, originally from Gazimulla-Tepa, Surkhandarya Province, Uzbekistan, Figure 1a) at a dedicated ancient DNA facility at the Department of Organismal Biology of Uppsala University. Prior to sampling, the bone was mechanically cleaned with a Dremel drill and a 3.0% (w/v) sodium hypochlorite solution, followed by UVed water. We extracted DNA following [14].

After DNA extraction, we prepared two double-stranded Illumina DNA sequencing libraries following [15]. Between each step the libraries were cleaned using Sera-Mag SPRI SpeedBeads (ThermoScientific) in 18% PEG-8000. Both libraries were sequenced independently at the National Genomics Infrastructure (NGI), SciLifeLab, Uppsala, on a NovaSeq 6000 Illumina DNA sequencer.

### Radiocarbon Dating

An additional 980 mg of bone fragment was obtained from the initial bone sample for radiocarbon dating. The direct radiocarbon dating was performed at the Tandem Laboratory, Uppsala University, and the obtained radiocarbon age was 2495±30.

The date was then calibrated by using the R package *rcarbon* [16] with atmospheric data from [17] (Figure 2).

**Figure 2:**
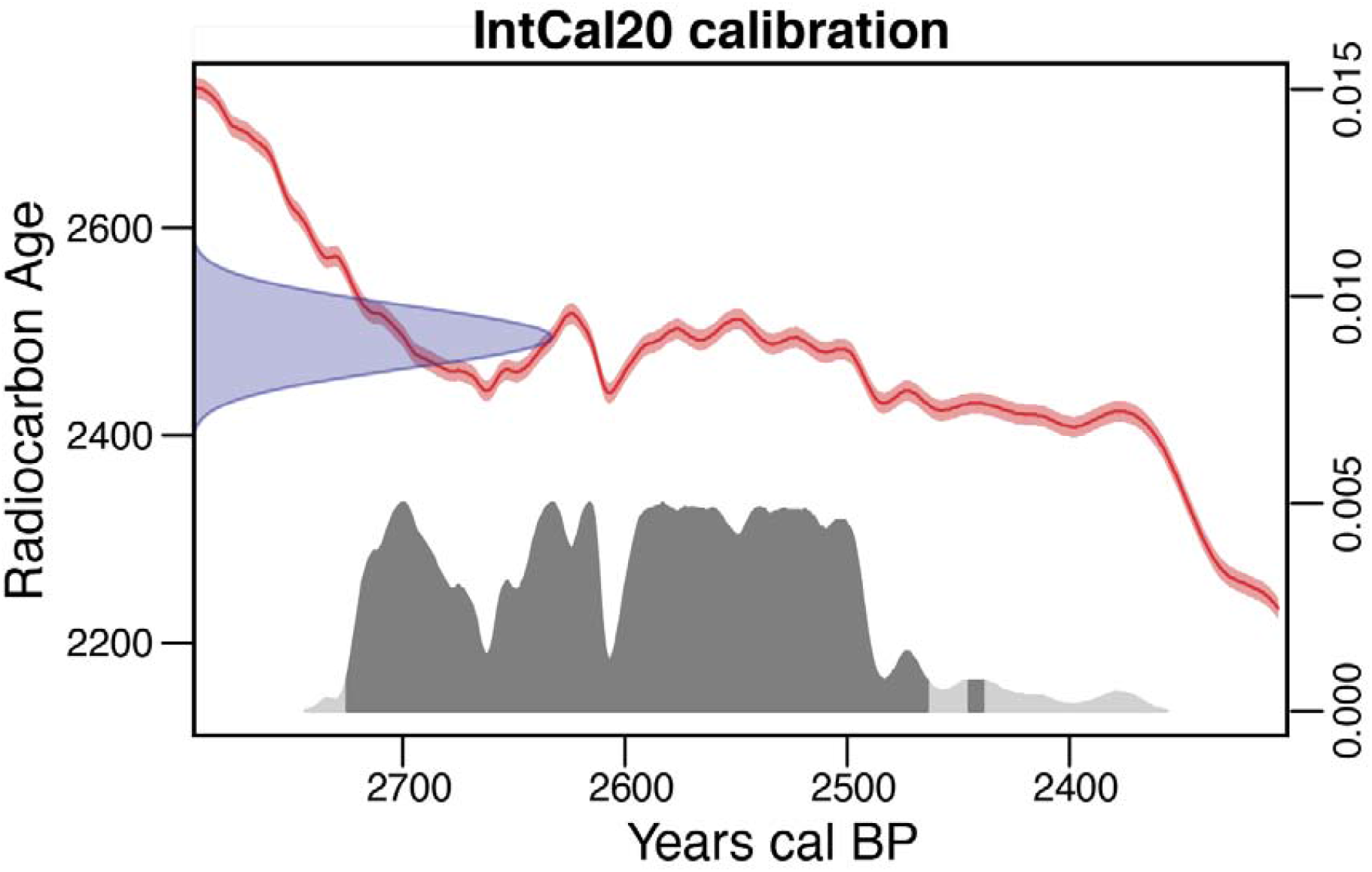
Calibrated radiocarbon age of AGAZ005 using *rcarbon* [16] and atmospheric data from IntCal20 [17].

### Data processing and authentication

We processed the resulting sequencing data by removing sequencing

adapters with CutAdapt [18], and merging the paired-end reads using Flash [19], requiring a minimum overlap of 11 bp and a maximum overlap of 150 bp. Due to the lack of a reference genome for the *Gazelle* genus we mapped the merged reads to the sheep genome Oar_v3.1 (GenBank assembly accession: GCA_000298735.1) in order to assess nucleotide misincorporation and damage patterns. We mapped the merged reads to the sheep reference genome using BWA aln parameters established for ancient DNA studies (“-l 1024 -n 0.01 -o 2”) [20], and created sorted BAM files with samtools 1.12 [21]. We removed PCR duplicates with Picard’s MarkDuplicate [22]. We also checked the biological sex of our sample using the Rx method [23] modified for a reference genome with 26 autosomes.

### Species identification and mitochondrial genome assembly

We first used MEGAN v.6.20.19 [24] to taxonomically assign the sequenced reads. Furthermore, we assembled the mitochondrial genome by mapping the reads to different mitochondrial reference genomes (Table 1) using MIA (https://github.com/mpieva/mapping-iterative-assembler). MIA is an iterative assembler designed to be used with ancient DNA, improving alignment and consensus on chemically damaged DNA. The final consensus sequence for each assembly was called with each base having a minimum of 3-fold depth coverage and two-thirds base agreement.

**Table 1:** Sequencing information for both libraries. Mapped reads to the nuclear sheep genome, since there is no gazelle genome currently available.

| Sample | Read Pairs | Merged Reads | Mapped Reads | % Mapped | % Duplicates |
| --- | --- | --- | --- | --- | --- |
| AGAZ005 (lib1) | 34,991,835 | 34,327,734 | 1,896,442 | 5.52% | 58% |
| AGAZ005 (lib2) | 21,264,728 | 21,204,583 | 514,280 | 2.59% | 43% |

### Phylogenetic analysis

In order to investigate this newly assembled mitogenome, we downloaded an additional 66 complete mitochondrial genomes from NCBI from all major Bovidae clades (Supplementary Table 1). We aligned all 67 mitogenomes using MAFFT [25] and inspected the alignment in SeaView 5.0.4 [26]. We reconstructed its phylogenetic tree through a Bayesian framework in MrBayes 3.2.7 [27], and also by using a Maximum Likelihood approach in IQ-Tree [28,29]. We used a GTR+G nucleotide substitution model in both analyses [30,31]. In MrBayes we ran three independent chains for 2 million generations ensuring a split frequency standard deviation below 0.01. In IQ-Tree we ran 1,000 bootstrap replicates (Supplementary Figure 2).

## Results

We sequenced about 56 million read pairs from two libraries. Approximately 4.3% of the merged reads mapped to the sheep reference genome (Table 1). The DNA sequences show fragmentation and deamination patterns characteristic for ancient DNA damages [32] indicating authenticity of the sequence data (Figure 1b and c). The sample was directly radiocarbon dated to 2725-2439 calBP (95.4% HDPI, Figure 2) confirming its Iron Age origin.

### Species Identification

Initially, we attempted to identify the species of AGAZ005 by using the taxonomic assignment of sequences implemented in MEGAN [24]. Due to the lack of available Antilopinae sequencing data, BLAST-based taxon assignment indicates only the presence of Bovidae sequences, but does not allow for precise species identification (Supplementary Figure 1). Since identification was not possible through this metagenomics approach, and considering this bone was found among several sheep bones [8], we mapped the merged reads to the sheep (*Ovis aries*) mitochondrial genome reference sequence NC_001941.1 using MIA (https://github.com/mpieva/mapping-iterative-assembler). Upon initial inspection we found large stretches of poorly assembled regions of the mitochondrion, suggesting that the sequences do not originate from a sheep. We then performed a BLAST search (blastn) [33] using the partially assembled mitochondrial sequences. This allowed us to identify the fragmented assembly as goitered gazelle (*Gazella subgutturosa*, 97% identity, Evalue = 1.9e-17). We identified this individual as a female, according to the results from the Rx method adapted from [23] (Rx = 0.818, 95% CI: 0.8027–0.8345), which is based on the number of sequencing reads mapping to the sheep X chromosome.

### Mitochondrial Genome Assembly

In order to assemble an improved mitochondrial genome for this specimen we adopted a strategy that would also further strengthen our taxonomic identification results. We mapped the merged reads to multiple Bovidae species using MIA, as described above. We observed the number of mapping reads, and also how many MIA iterations were necessary for final convergence for the assembly (Table 2). The final assembly using the *G. subgutturosa* reference sequence not only had the highest number of mapping reads, but also required the least amount of iterative mapping to converge.

**Table 2:** Mitochondrial genome assemblies statistics.

| Species | Common name | Mapped Reads | Average Coverage | Mapping iterations | Accession Number |
| --- | --- | --- | --- | --- | --- |
| ***Gazella subgutturosa*** | Goitered gazelle | **4500** | **23.212** | 4 | JN632644.1 |
| *Eudorcas rufifrons* | Red-fronted gazelle | 4171 | 21.896 | 30 | MG603682.1 |
| *Ovis aries* | Sheep | 3726 | 19.891 | 30 | MW364895.1 |
| *Rupicapra rupicapra* | Chamois | 3681 | 19.798 | 30 | FJ207539 |
| *Cephalophus dorsalis* | Bay duiker | 3589 | 19.356 | 30 | JN632615.1 |
| *Pseudois nayaur* | Bharal | 3553 | 18.830 | 30 | JX101653.1 |
| *Damaliscus pygargus* | Bontebok | 3487 | 18.976 | 30 | FJ207530.1 |
| *Oryx dammah* | Scimitar oryx | 3458 | 18.337 | 30 | JN869311.1 |
| *Capra ibex* | Alpine ibex | n/a | n/a | did not converge | FJ207526.1 |
| *Nanger dama* | Dama gazelle | n/a | n/a | did not converge | JN632665.1 |

### Phylogenetic placement

We built a phylogenetic tree under a Bayesian framework to investigate the relationship between our ancient sample and modern gazelle specimens (Figure 3). This result also further solidifies the finding that this bone fragment belongs to a *Gazella sp*. individual. The phylogenetic tree indicates that our sample is more closely related to the modern *Gazella sp*. individuals from China and Turkmenistan, including the subspecies *Gazella subgutturosa reginae* [34], when compared to the sand gazelle (*G. marica*, [35]). Unfortunately, the present tree is unable to resolve the position of Cephalophini in relation to the gazelles. While the placement of Raphicerina and Procaprina in relation to Antilopina agrees with [36], support values are relatively low and disagrees with the results from the Maximum Likelihood tree (Supplementary Figure 2).

**Figure 3:**
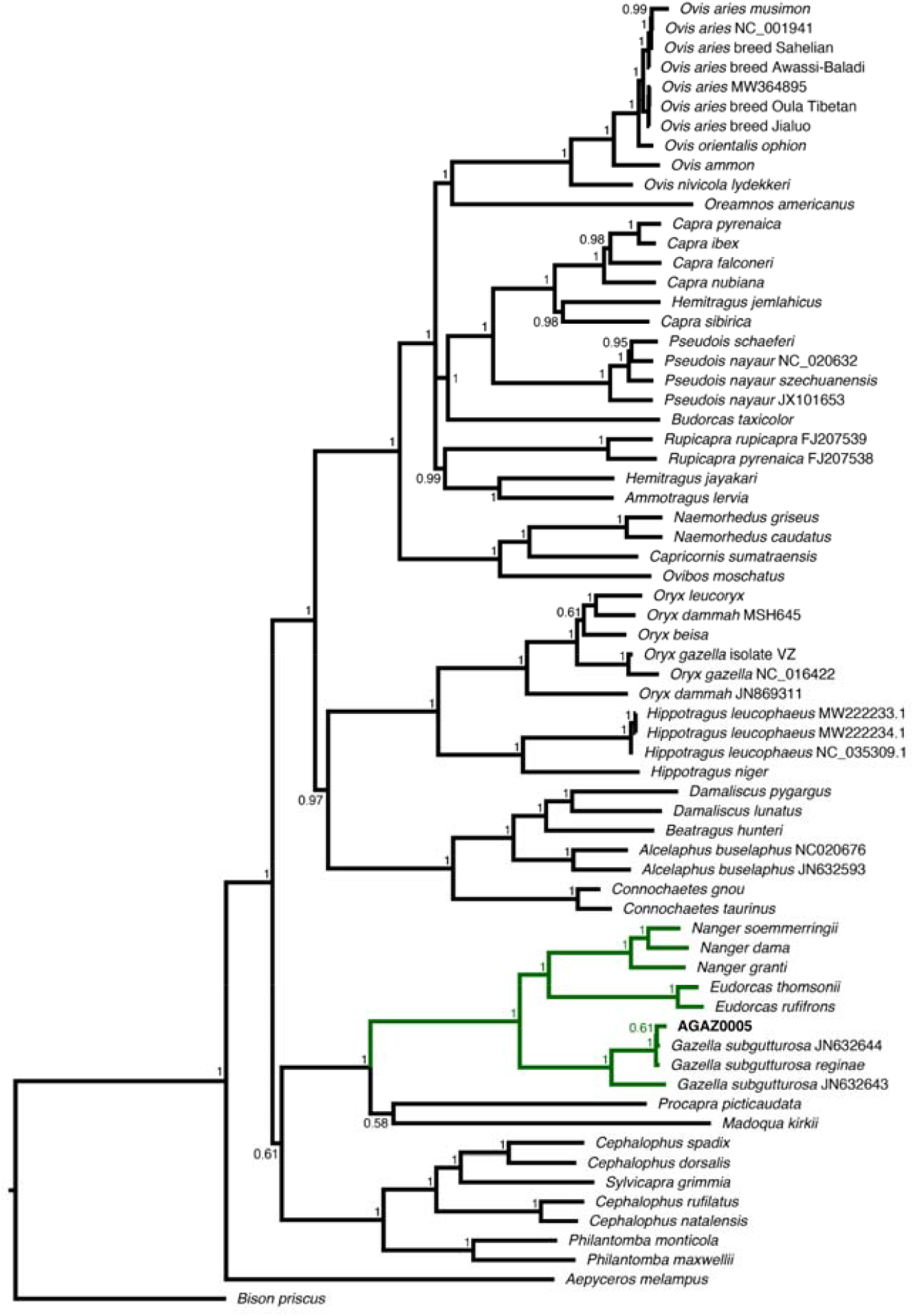
Bayesian inference phylogenetic tree for 68 mitochondrial genomes for a broad range of Bovidae species. In bold, our sample. In green the clade with all gazelle species represented in this tree.

## Discussion

Our study represents a direct genetic confirmation that goitered gazelles have been hunted in Iron age Uzbekistan around 2725-2439 calBP (95.4% HDPI). This is consistent with morphological analyses of the faunal remains found in the region which lists goitered gazelles as most common wild species [8]. Furthermore, a recent study using Zooarchaeology by mass spectrometry (ZooMS, [37]) on faunal remains from Neolithic and Early Bronze Age Kyrgyzstan also classified some specimen found at human settlements as “deer/*Saiga*/gazelle” [38].

Goitered gazelles have been subject to human hunting for several millennia across its entire range on the Asian continent [8,13,39]. As a result of habitat loss in combination with hunting and poaching, the species has been declining and today it is classified as “vulnerable” by the IUCN. Our study takes a first step towards genomic and temporal studies of this wide-ranging ungulate. The lack of modern genome-wide data for goitered gazelles restricted the types of analysis we were able to perform. Our results indicate that this ancestral population is closely related to modern individuals that inhabit Turkmenistan and the Qaidam basin in China, but is unable to determine which subspecies of *G. subgutturosa* it belongs to. The structure and potential gene flow between subspecies as well as the presumed recent population decline, however, would make the goitered gazelle an interesting subject for broad geographic sampling and population genomic studies. Smaller-scale studies based on microsatellite markers have already suggested local population structure [40]. Temporal genomic data could substantially enhance such studies of long-term processes. The genomic sequences generated for this project can be directly assigned to a point in time and space, and should therefore facilitate future studies of the population history of *G. subgutturosa*.

## Acknowledgements

We thank everyone involved in the excavations for their contributions.

## Authors’ contributions

A.E.R.S., O.S. and T.G. designed the research; A.E.R.S. performed the research; N. Bo. and L.S. conducted the archaeological excavations and here describes the chronological context; N. Bo., O.S., L.S. and N. Be. contributed samples; A.E.R.S. and T.G. analyzed data; and A.E.R.S., N.B., and T.G. wrote the paper with input from all authors.

## Data accessibility

Raw sequencing reads are available from the European Nucleotide Archive under accession PRJEB46820. The mitogenome for AGAZ005 (*Gazella subgutturosa*) was deposited at GenBank, accession number XXXX.

