## Supplementary Material for "Ancient DNA from a 2,700-year-old goitered gazelle (*Gazella subgutturosa*) confirms gazelle hunting in Iron Age Central Asia"

**Supplementary Table 1:** All mitochondrial genomes used in the phylogenetic analyses.

| **Species** | **Accession Number** |
| --- | --- |
| *Eudorcas thomsonii* | MG603682.1 |
| *Eudorcas rufifrons* | JN632634.1 |
| *Nanger granti* | JN632666.1 |
| *Nanger soemmerringii* | JN632667.1 |
| *Nanger dama* | JN632665.1 |
| *Gazella subgutturosa* | JN632644 |
| *Gazella subgutturosa reginae* | MW285638 |
| *Gazella subgutturosa* | JN632643 |
| *Procapra picticaudata* | MH345727.1 |
| *Madoqua kirkii* | JN632654.1 |
| *Aepyceros melampus* | JN632592.1 |
| *Oryx dammah* | JN869311.1 |
| *Oryx gazella* | JN632678.1 |
| *Oryx gazella* | NC_016422.1 |
| *Oryx beisa* | NC_020793.1 |
| *Oryx leucoryx* | NC_020732.1 |
| *Oryx dammah* voucher MSH645 | MT248297.1 |
| *Hippotragus niger variani* | KM245339.1 |
| *Hippotragus leucophaeus* | NC_035309.1 |
| *Hippotragus leucophaeus* | MW222233.1 |
| *Hippotragus leucophaeus* | MW222234.1 |
| *Connochaetes gnou* | JN632626.1 |
| *Connochaetes taurinus* | JN632628.1 |
| *Alcelaphus buselaphus* | NC_020676.1 |
| *Alcelaphus buselaphus* | JN632593.1 |
| *Beatragus hunteri* | NC_023542.1 |
| *Damaliscus pygargus* | FJ207530.1 |
| *Damaliscus lunatus* | KF955546.1 |
| *Ovibos moschatus* | FJ207536.1 |
| *Capricornis sumatraensis* | NC_020629.1 |
| *Naemorhedus griseus* | FJ207532.1 |
| *Naemorhedus caudatus* | NC_013751.1 |
| *Oreamnos americanus* | FJ207535.1 |
| *Budorcas taxicolor* | FJ207524.1 |
| *Rupicapra rupicapra* | FJ207539.1 |
| *Rupicapra pyrenaica* | FJ207538.1 |
| *Hemitragus jayakari* | FJ207523.1 |
| *Ammotragus lervia* | FJ207522.1 |
| *Pseudois nayaur* | JX101653.1 |
| *Pseudois schaeferi* | MK087727.1 |
| *Pseudois nayaur* | NC_020632.1 |
| *Pseudois nayaur szechuanensis* | KJ784494.1 |
| *Hemitragus jemlahicus* | FJ207531.1 |
| *Capra sibirica* | FJ207529.1 |
| *Capra nubiana* | FJ207527.1 |
| *Capra falconeri* | FJ207525.1 |
| *Capra pyrenaica* | FJ207528.1 |
| *Capra ibex* | FJ207526.1 |
| *Ovis nivicola lydekkeri* | MH779626.1 |
| *Ovis ammon ammon* | MN564883.1 |
| *Ovis orientalis ophion* | KF312238.2 |
| *Ovis aries musimon* | MG489885.1 |
| *Ovis aries* breed Jialuo | MK829158.1 |
| *Ovis aries* | MW364895.1 |
| *Ovis aries* breed Oula Tibetan | KU575248.1 |
| *Ovis aries* breed Awassi-Baladi | MW260509.1 |
| *Ovis aries* breed Sahelian | KF977846.1 |
| *Ovis aries* | NC_001941.1 |
| *Philantomba monticola* | JN632686.1 |
| *Philantomba maxwellii* | NC_020735.1 |
| *Cephalophus rufilatus* | JN632621.1 |
| *Cephalophus natalensis* | JN632618.1 |
| *Sylvicapra grimmia* | JN632701.1 |
| *Cephalophus spadix* | JN632623.1 |
| *Cephalophus dorsalis* | JN632615.1 |
| *Bison priscus* F3246 AE061 | KR350472.1 |

**Supplementary Figure 1:** BLAST-based taxon assignment of sequencing reads for both sequencing libraries as obtained in MEGAN.

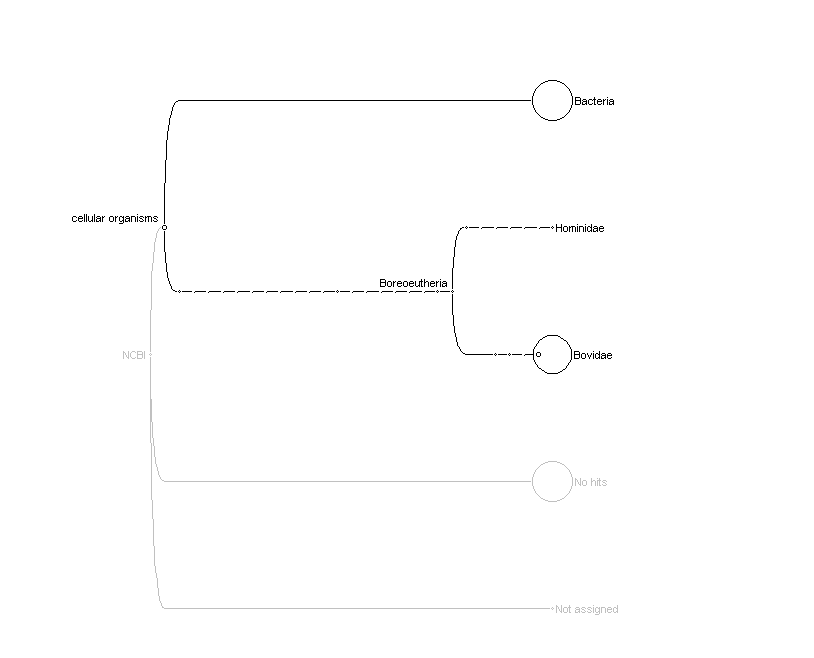

**Supplementary Figure 2:** Phylogenetic tree based on a Maximum Likelihood method, as implemented in IQ-Tree, under a GTR+G nucleotide substitution model and 1,000 bootstrap replicates.

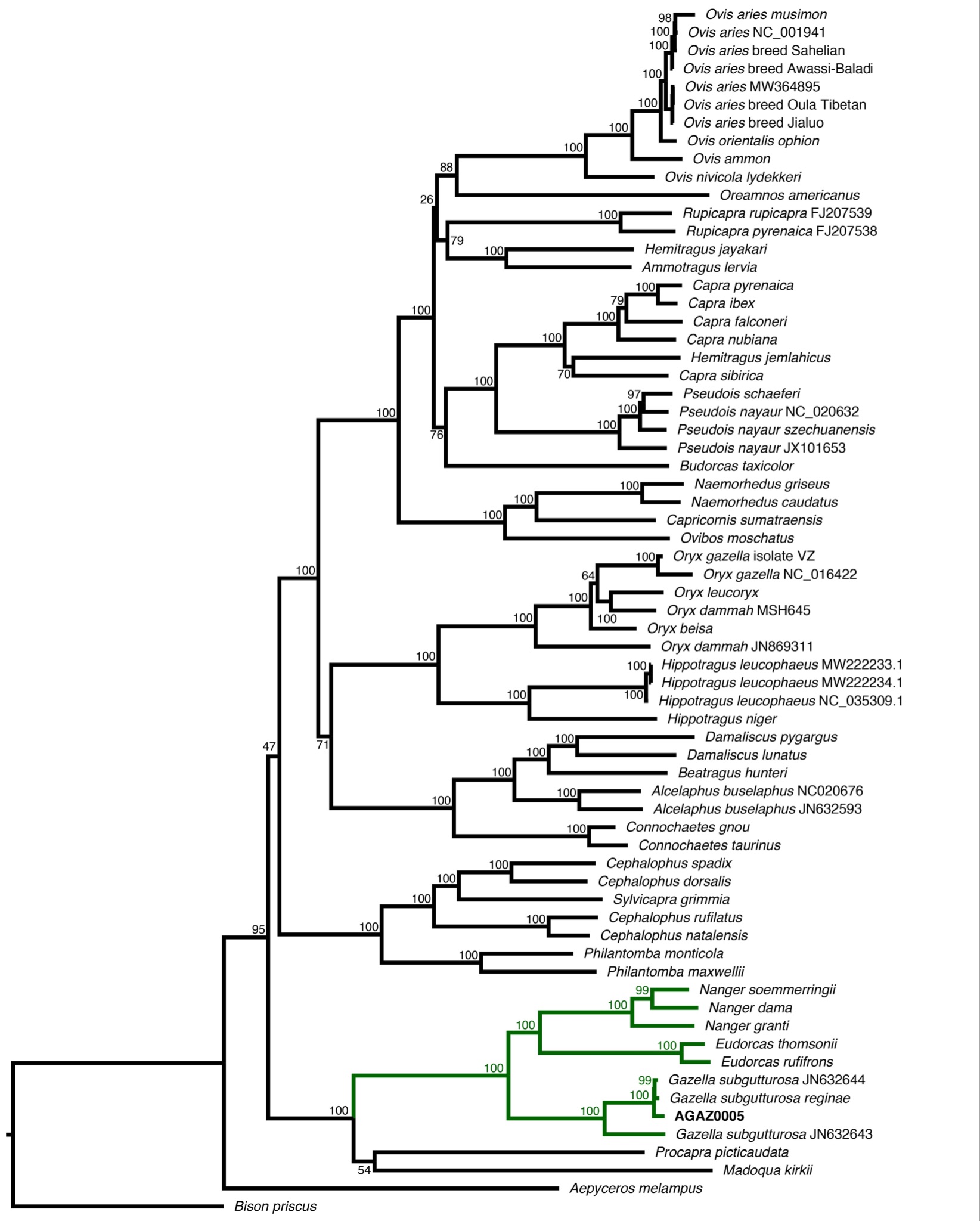
